# *In vitro* phenotypic susceptibility of HIV-1 non-group M to CCR5 inhibitor (Maraviroc): TROPI-CO study

**DOI:** 10.1101/2023.10.20.563343

**Authors:** Ségolène Gracias, Ikrame El Yaalaoui, Benoît Visseaux, Charlotte Charpentier, Diane Descamps, Charlène Martin, Fanny Lermechain, Jean-Christophe Plantier, Elodie Alessandri-Gradt

**Affiliations:** Univ Rouen Normandie, Univ Caen Normandie, INSERM, DYNAMICURE UMR 1311, CHU Rouen, Department of virology, F-76000 Rouen, France; Service de virologie, Université Paris Cité, INSERM, IAME, UMR 1137, AP-HP, Hôpital Bichat-Claude Bernard, Paris, France

**Author notes:** First author, Institut Pasteur, Université de Paris Cité, CNRS UMR3569, Virus Sensing and Signaling Unit, Paris, France. Second author, third author Laboratoire Cerba, Saint Ouen L’Aumône, France, Sixth author Institut Toulousain des Maladies Infectieuses et Inflammatoires (Infinity), Université de Toulouse, INSERM, CNRS, UPS, Toulouse, France.

## Abstract

The susceptibility of genetically divergent HIV-1 strains (HIV-1 non-M) from group O, N and P to the CCR5 co-receptor antagonist, Maraviroc (MVC) was investigated among a large panel of 45 clinical strains, representative of the genetic diversity. The results were compared to reference strains of HIV-1 group M (HIV-1/M) with known tropism. Among the non-M strains, a wide range of phenotypic susceptibilities to MVC was observed. The large majority of HIV-1/O strains (40/42) displayed a high susceptibility to MVC with median and mean IC_50_ values at 1.23 and 1.33 nM respectively, similar to the HIV-1/M R5 strain (1.89 nM). However, the 2 remaining HIV-1/O strains exhibited a lower susceptibility (IC_50_ at 482 and 496 nM), in accordance with their dual/mixed (DM) tropism. Interestingly, the 2 HIV-1/N strains demonstrated varying susceptibility patterns, despite always having relatively low IC_50_ values (2.87 and 47.5 nM). This emphasized the complexity of determining susceptibility solely based on IC_50_ values.

Our study examined the susceptibility of all HIV-1 non-M groups to MVC and correlated these findings with virus tropism (X4, R5 or DM). The results confirm the critical significance of tropism determination before initiating MVC treatment in patients infected with HIV-1 non-M. Furthermore, we advocate for the consideration of additional parameters, such as the slope of inhibition curves, to provide a more thorough characterization of phenotypic susceptibility profiles.

## INTRODUCTION

The high genetic diversity of HIV-1 has led to the current classification into four groups (M, N, O and P), but only HIV-1 group M (HIV-1/M) is pandemic (1). HIV-1 non-M group (HIV-1 non-M) are endemic in west-central Africa. HIV-1 group O (HIV-1/O) are subdivided into two main subgroups (T and H) (2) and are particularly found in Cameroon where it represents from 0.6% to 1% of HIV diagnoses. Only sporadic cases have been detected in other continents (3). HIV-1 group N (HIV-1/N) and HIV-1 group P (HIV-1/P) are even rarer, with respectively fewer than 20 and 2 reported cases of infection (3, 4). The french national survey network (RES-O) takes a census of all HIV-1/non-M infections in France and has already identified more than 140 cases of HIV-1/O infection, one primary HIV-1/N infection and one HIV-1/P infection, the most recent prototype strain of group P known to date.

The specific genetic diversity of the HIV-1 non-M viruses has known impacts on the antiretroviral treatments’ susceptibiliy. It has been previously demonstrated that the HIV-1/O were naturally resistant to the non-nucleoside reverse transcriptase inhibitors (NNRTI), particularly due to the Y181C resistance mutation present in 75% of the group O strains. The other antiretroviral agents have various *in vitro* efficacy, demonstrating the need of studying large panels of strains for HIV-1 non-M (5–7). The envelop region of HIV-1 non-M has a particularly high level of genetic diversity by comparison with HIV-1/M envelop amino acids sequences (8). The genetic sequence of the envelop V3 loop domain is commonly used as a predictor of the coreceptor use: CXCR4, CCR5 or both ; defining the virus tropism, usually called X4, R5 or dual/mixed (DM) respectively (9–11). Maraviroc (MVC) is actually the only CCR5 competitive antagonist commercialized since 2007 for patient with treatment failure (12). Because of its mechanism of action, this molecule is only effective on R5 viruses (13). Even if R5 tropism are the most frequent in the population (14), a predictive assay of virus tropism must always be performed before MVC initiation (15). To date, MVC is an alternative therapeutic strategy, for the patients previously treated and who may have acquired multiple resistance to diverse antiretrovirals. In the context of a more limited therapeutic arsenal for HIV-1/O, MVC could be a choice molecule to achieve a full active therapeutic combination for patients with additional resistance mutations. Due to less than 40% of similarity between group O and M V3 loops sequence, it has previously been demonstrated that usual HIV-1/M genotypic rules failed to correctly predict the HIV-1 non-M tropism. There is only one study reporting the absence of correlation between 5 genotypic tools and *in vitro* phenotypic tropism prediction of 18 group O strains (11).

Considering the absence of HIV-1 non-M approved tropism assay prior to MVC administration and the lack of *in vitro* data about MVC efficiency on HIV-1 non-M strains, the aim of this study was to define the phenotypic susceptibility of a large HIV-1 non-M panel (45 strains) to MVC and correlates these findings to the phenotypic tropism obtained by an in-house fluorescent cell assay.

## RESULTS

We analyzed 45 HIV-1 non-M clinical isolates: 42 HIV-1/O, 2 HIV-1/N and 1 HIV-1/P representative of HIV-1 genetic diversity as demonstrated with phylogenetic tree based on V3 sequences (Fig. 1), by comparison to 4 HIV-1/M control strains.

**FIG 1.**
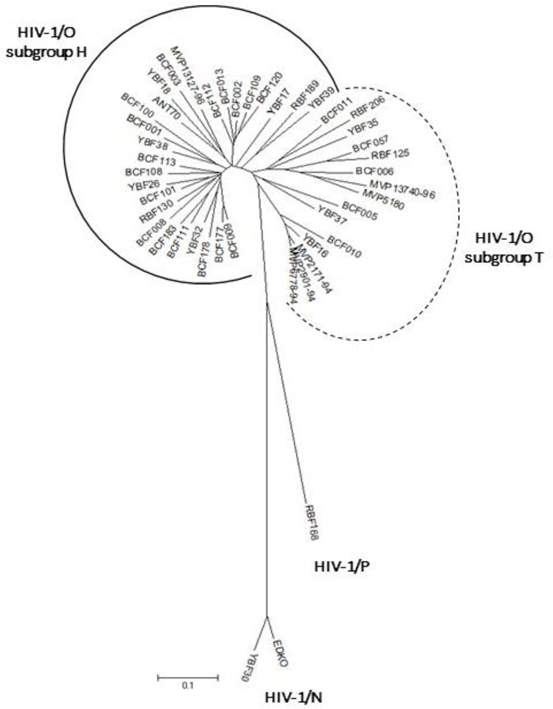
Phylogenetic tree of the 42 HIV-1/O, 2 HIV-1/N and 1 HIV-1/P V3 sequences from the strains included in this study

The HIV-1/M R5 strain (ARP1102) entry was efficiently blocked by MVC with an IC_50_ at 1.89 nM and MPI at 90.3% (Maximal plateau of inhibition), whereas the X4 strains (BRU-HXB2, ARP1196, and JR001) were all resistant (IC_50_ >1000 nM) and MPI at 0%, in accordance with what was expected for these reference strains (Table 1). The HIV-1/M DM tropism strain (ARP1129) was also resistant to MVC (IC_50_ >1000 nM) and MPI at 6.9%. Those 5 strains were also used to validate our tropism determination method because the tropism was already known.

**Table 1.** Results of tropism, IC_50_, MPI and hill slope obtained for each control and clinical strains of the panel.

| Strain | Group | Subgroup | Tropism | IC<sub>50</sub> (nM) | MPI (%) | Hill slope | GenBank n° |
| --- | --- | --- | --- | --- | --- | --- | --- |
| BCF006 | VIH-1/O | T | DM | 496 | 65.5 | ND |  |
| MVP5180 | VIH-1/O | T | DM | 482 | 29.6 | ND |  |
| YBF18 | VIH-1/O | H | DM | 0.57 | 94.7 | 3.71 |  |
| N1FR2011 | VIH-1/N |  | R5 | 47.5 | 63.7 | 1.09 | JN572926 |
| RBF168 | VIH-1/P |  | R5 | 3.68 | 88.3 | 0.91 | GU111555 |
| BCF100 | VIH-1/O | H | R5 | 3.22 | 94.2 | 1.57 |  |
| BCF108 | VIH-1/O | H | R5 | 3.1 | 85.5 | 2.15 |  |
| BCF101 | VIH-1/O | H | R5 | 3.09 | 95.5 | 1.06 |  |
| MVP2901-94 | VIH-1/O | T | R5 | 3.08 | 95 | 1.43 |  |
| BCF111 | VIH-1/O | H | R5 | 3.05 | 93.4 | 1.52 |  |
| BCF112 | VIH-1/O | H | R5 | 2.93 | 92.7 | 1.05 |  |
| YBF30 | VIH-1/N |  | R5 | 2.87 | 94.3 | 1.1 | AJ006022 |
| BCF177 | VIH-1/O | H | R5 | 2.66 | 90.6 | 1.57 |  |
| BCF010 | VIH-1/O | T | R5 | 2.57 | 97.3 | 1.08 |  |
| BCF113 | VIH-1/O | H | R5 | 2.34 | 94.2 | 2.23 |  |
| BCF057 | VIH-1/O | T | R5 | 2.21 | 96.9 | 1.47 |  |
| MVP2171-94 | VIH-1/O | T | R5 | 1.99 | 95.5 | 1.64 |  |
| BCF011 | VIH-1/O | T | R5 | 1.96 | 94.2 | 1.14 |  |
| BCF001 | VIH-1/O | H | R5 | 1.63 | 96.6 | 1.3 |  |
| BCF013 | VIH-1/O | H | R5 | 1.61 | 97.9 | 1.83 |  |
| YBF32 | VIH-1/O | H | R5 | 1.38 | 95.7 | 0.77 |  |
| RBF130 | VIH-1/O | H | R5 | 1.3 | 96.4 | 1.21 |  |
| BCF003 | VIH-1/O | H | R5 | 1.28 | 95 | 0.9 |  |
| RBF206 | VIH-1/O | T | R5 | 1.28 | 97.3 | 0.68 |  |
| BCF120 | VIH-1/O | H | R5 | 1.27 | 95.4 | 1.37 |  |
| YBF38 | VIH-1/O | H | R5 | 1.26 | 93.9 | 2.12 |  |
| BCF009 | VIH-1/O | H | R5 | 1.2 | 96.9 | 0.94 |  |
| ANT70 | VIH-1/O | H | R5 | 1.09 | 95.6 | 1.89 |  |
| MVP13740-96 | VIH-1/O | T | R5 | 0.79 | 95.1 | 0.56 |  |
| YBF16 | VIH-1/O | T | R5 | 0.75 | 97.2 | 1.48 |  |
| BCF109 | VIH-1/O | H | R5 | 0.75 | 96.1 | 1.16 |  |
| BCF005 | VIH-1/O | T | R5 | 0.71 | 97.3 | 0.95 |  |
| MVP6778-94 | VIH-1/O | T | R5 | 0.67 | 98.1 | 2.32 |  |
| MVP13127-96 | VIH-1/O | H | R5 | 0.58 | 98 | 1.62 |  |
| BCF002 | VIH-1/O | H | R5 | 0.58 | 98.9 | 0.81 |  |
| RBF189 | VIH-1/O | H | R5 | 0.56 | 98.8 | 1.09 |  |
| BCF178 | VIH-1/O | H | R5 | 0.47 | 98 | 1.3 |  |
| BCF183 | VIH-1/O | H | R5 | 0.47 | 96.7 | 0.94 |  |
| RBF125 | VIH-1/O | T | R5 | 0.32 | 97.7 | 0.62 |  |
| YBF35 | VIH-1/O | T | R5 | 0.18 | 97.2 | 1.12 |  |
| YBF17 | VIH-1/O | H | R5 | 0.11 | 98 | 1.43 |  |
| BCF008 | VIH-1/O | H | R5 | 0.07 | 97.2 | 1.53 |  |
| YBF26 | VIH-1/O | H | R5 | 0.01 | 89.9 | 0.7 |  |
| YBF39 | VIH-1/O | H | R5 | 0.005 | 85.8 | 0.67 |  |
| YBF37 | VIH-1/O | T | R5 | 0.003 | 90.8 | 0.73 |  |

**Table 2.**
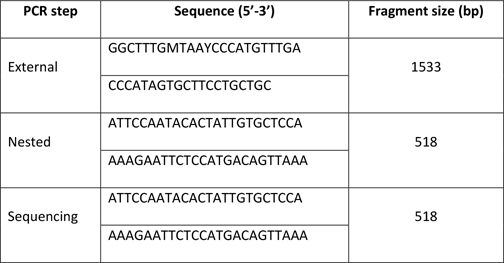
Primers used for C2V3 region of HIV-1/O amplification and sequencing.

Among the 45 available HIV-1 non-M strains, forty HIV-1/O were susceptible to MVC with IC_50_ between 0.003 (YBF37) and 3.22 nM (BCF100) and a corresponding median and mean IC_50_ at 1.23 and 1.33 nM respectively (Fig. 2). The mean MPI (min ; max) was at 95.3% (85.5 ; 98.9) (Table 1). The mean IC_50_ of these forty strains did not differ from the group M susceptible strain (non-significant). By our phenotypic tropism assay, all the strains demonstrated a R5 tropism except one (YBF18) displaying a DM tropism but with a high susceptibility to MVC: IC_50_ at 0.57 nM and 94.7% MPI.

**FIG 2.**
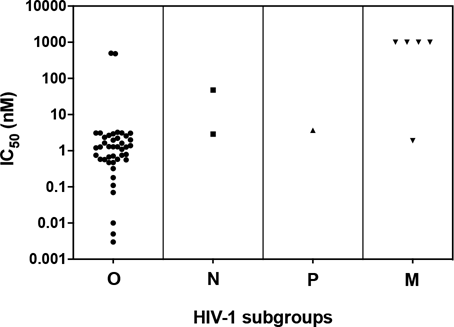
Comparison of HIV-1 non-M IC_50_ (nM) (42 HIV-1/O, 2 HIV-1/N and 1 HIV-1/P strains). The strains with a decrease of less than 1000 HIV GE/µL of viral load between the lowest and the highest concentration are considered as having a low susceptibility to MVC

No strain of the non-M panel showed a resistant profile similar to the X4 group M reference (Fig. 2). Although, the two remaining HIV-1/O strains expressed a very low MVC-sensitibe phenotype (MVP5180 and BCF006) with IC_50_ and MPI at 482 nM; 30% and 496 nM; 66% respectively. Those 2 strains showed DM profiles on the phenotypic tropism assay (Table 1) and have IC_50_ statistically different (p = 0.016) from the M reference R5 strain.

About the two HIV-1/N strains, the first one revealed a MVC-susceptibility with IC_50_ and MPI at 2.87 nM and 94% (YBF30). The second one (N1FR2011) was less susceptible to MVC with an IC_50_ at 47.5 nM and a MPI >50% (63.7%), whereas it has a R5 phenotypic tropism (Table 1). Despite this high difference of MVC-susceptibilities, the statistical analysis between the IC_50_ mean of the N group and the M reference strain remained non-significant (probably due to a lack of statistical power).

The HIV-1/P strain (RBF168) showed an IC_50_ at 3.68 nM and MPI at 88 %, this strain showed a R5 phenotypic tropism.

This wide range of various MVC-susceptibilities among the non-M panel is represented in Fig. 2 and 3. We distinguished the strains with low MPI (<50%) expressing weak susceptibility (N=1, MVP5180) and the strains with high susceptibility MPI>70% (N=40) representing the majority of the panel and the strains with intermediate susceptibility 50<MPI<70% (2 strains). Furthermore, we noticed among the highly susceptible strains, different behaviors from the shape of the inhibition curve. In this analyze, the two DM HIV-1/O strains less sensitive were not included. Thus, three groups of susceptible strains could be identified (Fig. 3) from the hill-slope value of the inhibition curve: the group with the steeper slopes [-4 ; -2[included 5 HIV-1/O strains with median IC_50_ (min ; max) 0.79 (0.57 ; 31). This group also included the DM HIV-1/O YBF18 MVC-susceptible strain. The group with intermediate slopes between [-2 ; -0.80[was the most important with 31 strains (20 subtypes H ; 8 subtypes T ; 2 HIV-1/N ; 1 HIV-1/P). A wide range of IC_50_ was also observed in this group with a median IC_50_ (min;max) at 1.28 nM (0.07 ; 47.5). Then the last group with slopes close to 0 [-0.8 ; -0.5[comprised 7 HIV-1/O with a corresponding median IC_50_ at 0.71 nM (0.03 ; 1.28).

**FIG 3.**
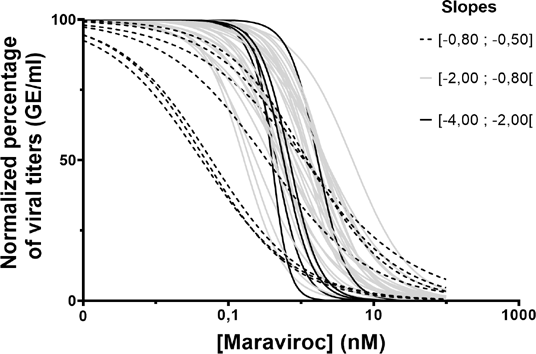
Susceptibility of 42 HIV-1/O expressed by normalized viral replication, on Maraviroc serial concentrations. Each curve depicts a different strain, the different patterns are used to group the strains with similar slopes.

The hypervariability of the C2V3 region across and within each HIV-1 non-M groups, made difficult the search of a genotypic correlation, but we noticed that the MVC susceptible DM HIV-1/O strain YBF18, had a non-conservative genomic pattern in the nine first amino-acid (MTCRRPA) (Fig. 4). There was no evident specific pattern or mutation associated with the highest fold change (FC), neither with the DM tropism. Comparing the genetic sequence of the three slopes groups, we noticed that 3 in 5 strains showing the highest slopes (<-2) harbored an insertion (Asparagine N our Alanine A) in position 7.

**FIG 4.**
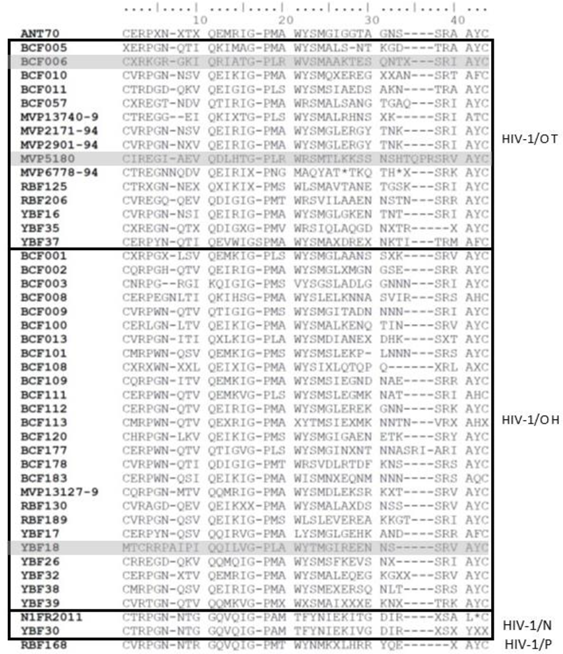
Alignment of V3 amino acid sequences from the HIV-1 non-M panel

## DISCUSSION

The aim of the TROPI-CO study was to explore the *in vitro* susceptibility of various HIV-1 non-M strains to MVC, to link these results to the R5, X4 or DM tropism, determined by a phenotypic assay and to explore a potential genotypic pattern associated to phenotypic tropism or MVC-susceptibility.

The MVC-susceptibilities for the HIV-1/M strains, were consistent to what was expected from the identified tropism, validating our cellular assays (for tropism and susceptibility determination). The large panel of HIV-1 non-M strains showed a global susceptibility to MVC, with a mean IC_50_ value at 1.23 nM, by comparison to group M (IC_50_ at 1.89 nM). Although a variability between culture methods can exist, these results are in accordance with the range of IC_50_ published by EMEA for O and M susceptible strains: to 8.9 nM (16). In a previous study conducted on 18 HIV-1/O strains, using a U87 human glioma cells based assay, MVC-susceptibility demonstrated a higher IC_50_ variability for HIV-1/O (ranging from 1 to 315 nM, mean at 51.2 nM) than for HIV-1/M (from 2 to 102 nM, mean at 32.4 nM) (17). In this *in vitro* study with non-primary cell line, HIV-1/O seems less susceptible to MVC. Fifteen strains were R5-tropic and 3 DM, but without mention about the corresponding IC_50_ values (17). When comparing to our panel, 12 HIV-1/O were also included in our study (29% strains in common strains). We can suppose that the higher IC_50_ came from the DM viruses, as what we observed from our panel: the two strains with FC around 400, were DM-tropic. Our observation also showed that there was no complete correlation between MVC susceptibility and R5 tropism, because one DM strain (YBF18) showed a full susceptibility to MVC (MPI at 94.7% and FC < 1). In fact, the literature has reported an inefficiency of MVC with this tropism (18, 19). It is possible that the DM tropism is not equally distributed among the strains sub-populations, with a higher proportion of R5 susceptible viruses in the YVF18 strain. This hypothesis still needs to be further explored by cell flow cytometric-directed analyzes for example.

We also noticed that the amino acid composition of the V3 loop of YBF18 is very singular, with the presence of two consecutive basic amino acid (R=Arginine) in the unique 9 amino-acid long-pattern (MTCRRPA).

The absence of MVC-resistant strain in our HIV-1 non-M panel is underlined by the absence of X4 tropism (IC_50_>1000 nM and MPI around 0%). As far as we know, there is no description of X4 HIV-1/O (or other non-M) strain in the literature (3, 17, 20), potentially meaning that the virus may use others coreceptors for the early steps of entry in the cell host. This could emphasize a potential role of an antiretroviral therapy including MVC, for those naturally highly divergent viruses. Regarding more deeply into the MVC-susceptibility, we noticed that a high IC_50_ value (and so a high FC) was not always associated with a weak MPI. As a consequence, those two elements cannot completely characterize the susceptibility profile of the strain. In addition, concerning the strains with MPI>90%, all were susceptible to MVC but presented different inhibition curve slopes, suggesting potential differences in therapeutic response to MVC. That is why we proposed to also considered the hill slope for additional parameters of susceptibility characterization as to IC_50_/FC and MPI. As an example, the HIV-1/N strain (N1FR2011) showed a FC at 40, but remained susceptible: its inhibition curve showed a maximum decrease on the viral load between 10 and 100 nM concentrations, with a hill slope at -1.09 and therefore cannot be assigned to a resistant strain despite its high FC and intermediate MPI. Thus, a potential different therapeutic response could be expected for the infected patients.

## MATERIALS AND METHODS

### Cells and virus stocks

Peripheral blood mononuclear cells (PBMCs) were isolated at the ‘Etablissement Français du Sang’ (EFS) from 3 different healthy donors to avoid unique donor bias on type cells effects and extracted using Ficoll-Plaque Plus solution (GE Healthcare). PBMCs were re-suspended in RPMI-1640 medium (Lonza Bioscience) supplemented with 10% FBS (PAA laboratories), 50 µg/mL gentamicine (Panpharma), 0.2 µg/mL phytohemagglutinin (PHA) and grown at 37°C with 5% CO_2_. After 72 h, PBMCs were activated with interleukin-2 (IL-2) 450 UI/mL (Proleukin, Novartis pharma) and polybrene 2 µg/mL (Sigma-Aldrich) during at least 2 h before infection.

GHOST cells expressing CCR5 or CXCR4 were provided by the NIBSC. Those human osteosarcoma cells constitutively express CD4 receptor and the HIV coreceptor CCR5 or CXCR4. They also contain, as a HIV infection infection reporter gene, a HIV-2 long terminal repeat (LTR) sequence linked to a green fluorescent protein (GFP) gene. HIV infection induces viral Tat protein production, activates the HIV-2 LTR promoter and generates GFP expression. Thus, HIV entry can be monitored by fluorescence intensity observation (17). The cells were cultured in DMEM with 10% FBS (PAA laboratories) and 1% penicillin and streptomycin (Sigma-Aldrich). In order to maintain the expression of the coreceptors, hygromycin 100 µg/mL (Invitrogen), puromycin 10 µg/mL (Sigma-Aldrich) and geneticine G418 500 µg/mL (Invitrogen) were added in the maintenance medium, as recommended.

HIV-1 non-M strains were isolated from HIV positive patients (cells or plasma samples) cultivated 3 weeks on PBMCs collected from healthy donors (RPMI, 10% FBS supplemented, gentamycin 50 µg/mL, at 37°C with 5% CO_2_). Half of the medium was replaced twice a week. During this period, the amount of virus was regularly quantified in the supernatant by measuring the activity of the reverse transcriptase (Lenti RT Cavidi kit, Cavidi). Once the peak of activity of the viral enzyme was reached, the culture was stopped, and aliquots of viral supernatant stored at -80°C.

Five HIV-1/M strains (BRU-HXB2, JR001, ARP1196, ARP1102 and ARP1129) with previous characterized tropism one HIV-1/M strain with R5, 3 with X4 tropism (ARP1196, BRU-HXB2, JR001) and one DM tropism (ARP1129) were added as controls. These five strains were chosen as references for both drug phenotypic susceptibility and tropism assays. For this study, we analyzed 45 HIV-1 non-M clinical isolates: 42 HIV-1/O, 2 HIV-1/N and 1 HIV-1/P representative of HIV-1 genetic diversity as demonstrated with phylogenetic tree based on V3 sequences (Fig 1).

### Tropism assay

To determine the tropism of each strain, we used 2×10^4^ GHOST cells expressing co-receptor CCR5 or CXCR4 seeded in a 96-well plate and which were then incubated at 37°C, 5% CO_2_. After 12 h to obtain cells confluence, 75 µL of HIV supernatant was added in triplicates (final volume of 200 µL) during 3 h, then the cells were washed with PBS before adding 200 µL DMEM. At day 3, fluorescence from infected GHOST cells was checked by fluorescent microscopy to determine tropism. R5 tropism was characterized by fluorescent CCR5 GHOST cells (green) and non-fluorescent CXCR4 GHOST cells (colorless) and vice versa. If both cells turn out to be express a green fluorecence, a DM tropism was defined.

### Phenotypic assay and RNA extraction

The phenotypic assay was performed as previously described (21). Briefly fresh PBMCs PHA-stimulated were infected by 100 TCID_50_/mL of HIV-1 supernatant for 2 h and then cultivated with five serial dilution of increasing concentrations of MVC (0.1, 0.5, 2, 5, 10, and 100 nM) during 72 or 96 h. Each dilution was tested in quadruplicate in a 96-well plate. The quadruplicates were pooled and then RNA extraction was performed by the automated EZ1 advanced XL system (Qiagen) with EZ1 DSP Virus kit (Qiagen).

### qRT-PCR analyses

HIV-1/O RNA was quantified on a CFX96 Deep-Well (BioRad) by targeting integrase O thanks to primers and TaqMan probe: Forward TCTATTACAGAGACAGCAGAGAYC Reverse CTACTGCTCCYTCACCTTTCC and probe FAM-ACAGGAGYTGKGCCGGTCCTTTC Dark Quencher with RNA UltraSense™ One-Step Quantitative RT-PCR System (Invitrogen). The PCR was as followed: retro-transcription (RT) 15 min at 50°C, denaturation 2 min at 95°C followed by 50 cycles of cDNA amplification and denaturation 15 sec at 95°C and 30 sec at 60°C. Genome equivalent (GE) concentrations are determined by extrapolation from a standard curve generated from serial dilutions of total HIV-1/O RNA of a known concentration.

HIV-1/N and M strains were quantified by targeting the long terminal repeat (LTR) region using GENERIC HIV Charge virale kit (Biocentric) performed on CFX96 Deep-Well (Biorad) with the followed program RT 10 min at 50°C and 5 min at 95°C, amplification of cDNA 50 cycles 15 sec at 95°C, 1 min at 60°C.

HIV-1/P strain was quantified using Xpert HIV-1 viral load kit (Cepheid) that was proven as adapated to this strain from previous results of performance (22).

### Sequencing of the V3 loop region and phylogenetic analysis

Amplification of the C2V3 coding region of HIV-1/O was performed by PCR (Invitrogen Superscript III One step RT-PCR for long template kit). The PCR conditions were 30 min at 50°C, 2 min at 94°C then 35 cycles of amplification (30 sec at 94°C, 30 sec at 55°C, 2 min at 68°C) and finally 10 min at 68°C just before reaching 4°C for storage. A nested RT-PCR (QIAGEN hot start Taq master kit 1000 units) was then performed from 2 µL of previous amplicon for 15 min at 95°C then 35 cycles of amplification (30 sec at 94°C, 30 sec at 55°C and 90 sec at 72°C) to finish with 7 min at 72°C before storage at 4°C. The primers used are listed in Table 1. Sanger sequencing was performed on CEQ 8000 with the following program: 30 cycles (25 sec at 96°C, 25 sec at 50°C) followed with 4 min at 60°C and then 4°C using CEQ DTCS Quick Start kit. The sequences were analyzed and aligned using Genome Lab and MEGA 6 software to construct the phylogenetic tree by the *Neighbor-joining (500 bootstraps, calculation with the 2 parameters Kimura method)*. The V3 sequences for the 2 HIV-1/N and HIV-1/P strains were extracted from sequences previously published in the Genbank (JN572926, AJ006022 and GU111555 respectively).

### Statistical Analysis

The analysis of concentrations of viral RNA in GE/ml were based on a qRT-PCR standardization to report the concentration from the cycle threshold.

The drug concentration that inhibits strain replication by fifty percent (IC_50_) was defined. The maximum percentage of inhibition (MPI) corresponding to the point where the viral replication reaches its minimum under drugs effect, by comparison to the point without drug, was also define. These parameters were calculated with Microsoft office Excel software. Graph Pad Prism was used for drawing the graphs and calculate the hill slope after transformation of the concentration in decimal logarithm and normalization with 100% for the highest value and 0% for the lowest. Unpaired t-test (p<0.05) was applied on the mean of our IC_50_ of HIV-1/M and IC_50_ of HIV-1/M strain reference.

## Acknowledgements

The TROPI-CO project was supported by the Rouen University Hospital and the region of Rouen Normandy, and Santé Publique France. The funders had no role in study design, data collection and interpretation, or the decision to submit the work for publication.

We thank the IAME laboratory from Bichat Hospital and Université Paris Cité, for providing the drug. We thank Gatien Durand and Romain Legros for their contribution to the project.

